# Toxin structure shapes palatability in a chemically defended butterfly

**DOI:** 10.64898/2026.09.18.751997

**Authors:** Erik Todd, Aaron Denton, Dheeraj Halali, Glennis Julian, Adria LeBouef, Chris D. Jiggins, Erika C. Pinheiro de Castro

## Abstract

The toxicity of chemical defences is well studied, but the potential contribution of compound structure to predator deterrence remains largely unexplored. Whether predation acts more strongly on toxicity or unpalatability remains largely untested, partly because few systems allow toxin structure to vary independently of quantity. *Heliconius sara* larvae provide such a system: those reared on *Passiflora auriculata* sequester cyclopentenyl cyanogenic glucosides (CGs), while those reared on *P. biflora* biosynthesise comparable quantities of aliphatic CGs. Using two invertebrate predators, *Camponotus floridanus* ants and *Hierodula membranacea* mantids, we tested whether this structural difference affects palatability independent of toxicity. Mantids rejected larvae with cyclopentenyl CGs more often than larvae with aliphatic CGs, despite no detectable difference in total CG content. This pattern was mirrored in extract-based assays with ants, independently of cyanide release: extracts with cyclopentenyl CGs remained deterrent, while extracts with aliphatic CGs did not differ in deterrence from water. Live larvae, by contrast, elicited similar responses from ants regardless of CG structure. These results show that variation in toxin structure can strongly affect palatability, with some compounds conferring greater protection than others. This demonstrates the importance of chemical structural diversity in the evolution of chemical defences.

## Introduction

Although often used synonymously, toxicity and unpalatability are distinct [1]. Toxicity directly reduces predator fitness, whereas unpalatability is a negative sensorial experience perceived via gustatory or olfactory cues [1]. Predators can learn from aversive encounters to avoid toxic or unpalatable prey in the future [2]. The effectiveness of chemical defences therefore depends not only on their toxicity, but also on how readily they are identified [2], [3], [4], [5], [6]. However, the question of whether predation acts more strongly on toxicity or unpalatability remains largely untested [7]. Furthermore, most studies investigating the palatability of chemically defended invertebrates have used avian predators [5], [8]. Different predator taxa may respond differently to diverse chemical defences [9]. A broad range of model predators is therefore needed to fully understand the role of predation in the evolution of toxicity and unpalatability of chemical defences. Understanding this ‘palatability spectrum’ is also important to resolve outstanding evolutionary questions, such as the importance of quasi-Batesian mimicry [10], [11].

Toxic animals, particularly insects, generally biosynthesise defensive compounds *de novo* or acquire them through dietary sequestration [12], [13], [14]. Biosynthesis, while rarer than sequestration [15], has evolved independently in several lineages, and allows toxicity to be maintained without dependence on a specific food source [13].

In several species, the same individual can either sequester or biosynthesise toxins, depending on the environmental context [14], [16]. As sequestration and biosynthesis rely on distinct biochemical pathways [15], this plasticity can generate variation in defensive compounds between conspecifics [16]. This offers a route to test how chemical structures influence chemical-defence effectiveness, as compound structure can be varied independently of toxicity.

The *Heliconius* butterflies, a diverse clade of toxic butterflies, provide a system for investigating this question. Several species exhibit plasticity in how they acquire cyanogenic glucosides (CGs) [16], their principal chemical defence [17]. Upon tissue disruption, CGs come into contact with hydrolytic enzymes and are broken down into aglycones and hydrogen cyanide (HCN) [18], [19]. This release is not instantaneous and can occur over several minutes following tissue damage [18], [20]. CGs vary structurally, for instance in their aglycone backbone [14], [21]. Variation in both aglycones and intact CG could be perceived by predators independently of cyanide release. The maintenance of variation in chemical defence strategies within the *Heliconius* butterflies, and its implications for fitness remain poorly understood, in part because the defensive properties of different CGs have not been previously studied. A comparable pattern has been demonstrated in milkweed bugs, where sequestered cardenolide structure, rather than quantity, determined protection against a specific invertebrate predator [22].

Here, we test whether structural variation between CG classes affects the palatability of *Heliconius sara* larvae. In *H. sara*, larval diet influences toxin acquisition strategy, which in turn leads to differences in the chemical structures of CGs. This allows hostplant availability to be manipulated to generate structurally distinct chemical phenotypes, despite comparable total CG content. Here we combine live-larva assays using two invertebrate predators, *Camponotus floridanus* (carpenter ants) and *Hierodula membranacea* (giant Asian mantids), with extract-based assays in which intact CG extracts are painted onto a palatable control species. Together, these approaches allow us to investigate the effect of compound structure on predator responses independent of overall toxin levels. We can therefore directly explore the role of unpalatability in chemical defences.

## Methods

### Study systems and rearing conditions

An insectary population of *H. sara*, reared in the Department of Zoology, University of Cambridge, was maintained under controlled conditions (25°C, 75% RH, 12h light:dark cycle) and used throughout this study. Egg clusters were transferred to rearing cages (100cm x 40cm x 40cm) and raised on either *P. biflora*, on which larvae biosynthesise aliphatic CG (linamarin and lotaustralin), or *P. auriculata*, on which larvae predominantly sequester cyclopentenyl CG (epivolkenin) [14], [21]. Larvae were fed *ad libitum* with fresh cuttings of their assigned host plant until the third to fourth instar, when they were used as prey in unpalatability assays. *Bicyclus anynana*, reared on maize, and *Galleria mellonella*, sourced from online suppliers, were used as non-toxic control species across experiments.

Young *H. membranacea* nymphs were sourced from an online supplier (Mantis House Ltd. UK) for use as predators in unpalatability assays. Mantids were kept individually in plastic tubes (500ml) under the same controlled conditions as above and fed *Drosophila hydei* every other day, sourced from a commercial live-food supplier. Fifth and sixth instar nymphs were used in the assays. Mantids are known natural predators of butterflies [23] and have been used as predator models in previous unpalatability assays [24], [25].

*Camponotus floridanus* workers were also used as predators, drawn from an insectary population maintained in the Department of Zoology. Colonies were housed in 5L buckets under controlled laboratory conditions and fed once weekly. Ants regularly visit extrafloral nectaries on Passiflora plants [26], [27] and consequently have frequent contact with *Heliconius* larvae in the wild [28], making them ecologically relevant predators in this system.

### Experiment 1: Palatability assays using live larvae and mantid predators

We collected clutches of *H. sara* eggs and separated them into two dietary groups, reared in separate enclosures. Larvae were provided fresh young stems twice weekly from either *P. auriculata* or *P. biflora*. Larvae raised on each diet did not differ significantly in total CG content (see Results) but differed in CG composition. Larval stage was matched across all diets and species at the point of testing.

Trials were conducted in 20 x 20 x 20cm Perspex enclosures fitted with a diagonal perch. Mantids were starved for 48h and acclimated in the testing enclosure for 5 minutes before each trial. A single larva was introduced to mark the start of each 15-minute trial. Rejection was recorded as a binary outcome, defined as a mantid actively discarding a prey item during the trial. Face wiping, an established response to aversive food where mantids wipe their mandibles on the substrate [29], [30] was recorded as a second binary response variable. Four larval types were tested: *H. sara* raised on *P. auriculata* (n = 20), *H. sara* raised on *P. biflora* (n = 23), wax moth controls (n = 26), and *B. anynana* (n = 14). Twenty individual mantids were used across these trials, each tested multiple times; mantid identity was tested as a random effect in the statistical model (see Statistical analysis).

Larvae from each brood and treatment group used during testing were frozen at −80°C. Total CG content was quantified to determine whether toxicity differed between experimental groups. LC-MS/MS was used to quantify cyanogenic glucoside content, following the protocol outlined in [14]. Frozen larvae were homogenised in 80% methanol, centrifuged at 10,000g for 5 minutes, and the supernatant analysed using LC-MS/MS (Agilent 1100 Series LC; Bruker HCT-Ultra ion trap mass spectrometer). Total glucoside content was estimated from extracted ion chromatogram peak areas and quantified using calibration curves.

### Experiment 2: Palatability assays using live larvae and ant predators

*H. sara* larvae were reared as described above; third instar larvae were used for testing throughout. Individual *H. sara* larvae, with either aliphatic or cyclopentenyl CG, were presented to single *C. floridanus* workers in Sterilin 50mm Petri dishes. Control trials used wax moth larvae. Ants were habituated in the dishes for 5 minutes prior to testing, and individual trials lasted 10 minutes. Batches of 9 trials were filmed concurrently from above and bottom-lit with an LED light. Dishes were cleaned with ethanol between trials. As larvae had to be used upon reaching the desired instar, treatment groups could not always be tested on the same day; of 89 trials conducted, 36 used *H. sara* raised on *P. auriculata*, 27 used *H. sara* raised on *P. biflora*, and 26 used wax moth larvae.

Trials were filmed at 25 fps using a camera mounted above the arena. Custom Python tracking code based on [31] was used to identify frames in which ants were interacting with larvae [32]. Interactions were classified as events where ant and larvae contours overlapped. Single-frame interactions (0.04s) were excluded as likely tracking artefacts, given their excess frequency when the interaction-duration distribution was analysed. The total time ants spent interacting with larvae per trial was the response variable examined.

### Experiment 3: Palatability assays using cyanogenic glucoside extracts

The live larval assays cannot distinguish between physical and chemical sources of aversiveness. Furthermore, because tissue disruption during a real predation event releases hydrogen cyanide, they cannot determine whether an aversive response reflects the released toxin itself or properties of the intact CG. CG extracts were therefore prepared and applied to wax moth larvae prior to presentation to *C. floridanus* workers.

Extracts were prepared using *H. sara* larvae raised on *P. biflora* and *P. auriculata* (n = 13 for each diet), with all siblings being from the same brood and at the same developmental stage. Frozen larvae were macerated in 10mL of 80% methanol, centrifuged at 7500×g for 10 minutes, and the supernatant dried for 20 hours using a Savant DNA120 SpeedVac Concentrator. This extraction method is expected to denature β-glucosidase and substantially limit enzymatic hydrolysis of cyanogenic glucosides during processing [33]. The dried residue was resuspended in 5mL of water, aliquoted (400µL), and frozen at −20°C. On testing days, extracts were defrosted and kept on ice. Each wax moth larva received two 20µL applications of extract or water, pipetted in three dorsal drops and spread with a clean treatment-dedicated fine-tipped paintbrush. Once dry, larvae were presented to single *C. floridanus* workers in Sterilin 50mm Petri dishes, preceded by a 5-minute ant acclimation period. Trials lasted 15 minutes. Fluon was applied to dish sides to prevent ants from accessing the lid. A fresh ant and larva were used in each trial, and 54 trials were conducted per treatment group.

Filming, tracking, and interaction classification followed the protocol described for Experiment 2, and total interaction time was again the response variable examined.

### Statistical analysis

All analyses were conducted in R (version 4.3.3) [34]. Unless stated otherwise, treatment group was included as a fixed effect, and model diagnostics (DHARMa simulated residuals[35]) indicated no deviations from model assumptions in any model. All pairwise comparisons used Tukey correction via emmeans [36].

**Mantid rejection and face wiping**: rejection was modelled using a binomial GLM (logit link); face wiping was modelled using a binomial GLMM (logit link), fitted using glmmTMB [37], [38]. Random effects for mantid identity and testing day were tested for each; testing day was dropped from both (non-significant), while mantid identity was retained only for face wiping (significant), which is accordingly reported from the GLMM. Controls were excluded from both models due to complete separation from zero rejections. Larval weight, included as a scaled covariate, was not significant. Ant live-larva assay: total interaction time was modelled using a gamma GLM (log link). A batch term (colony × day) could not be tested due to trial-number constraints per combination.

**Ant extract assay**: total interaction time was modelled using a gamma GLM (log link). A batch term (colony × day) was tested, found non-significant, and excluded from the final model. Standard errors were nonetheless clustered by batch as a conservative measure, and all reported estimates and pairwise comparisons for this model use these batch-clustered robust standard errors [39], [40].

## Results

### Mantids reject butterflies with sequestered cyclopentenyl CGs more often than those with biosynthesised aliphatic CGs

We compared palatability of *H. sara* larvae with different CG profiles against mantid predators. *H. sara* larvae with cyclopentenyl CGs (sequestered) were rejected at a substantially higher rate than all other groups, with 12 of 20 trials resulting in rejection (60.0%), compared with 5 of 23 *H. sara* with aliphatic CGs (biosynthesised) (21.7%), 1 of 14 *B. anynana* (7.1%), and 0 of 26 wax moth controls (0%) (Figure 1a). The binomial model confirmed that *H. sara* larvae with cyclopentenyl CGs were significantly more likely to be rejected than larvae with aliphatic CGs (estimate = 1.87, SE = 0.712, z = 2.624, p = 0.0236) and *B. anynana* (estimate = 3.24, SE = 1.150, z = 2.809, p = 0.0138). *H. sara* with aliphatic CGs and *B. anynana* did not differ significantly in rejection probability (estimate = 1.37, SE = 1.160, z = 1.184, p = 0.463). Target-metabolomics revealed that diet did not affect the total CG content of *H. sara* (Wilcoxon rank-sum test: W = 26, p = 0.230), just their chemical profile.

**Figure 1:**
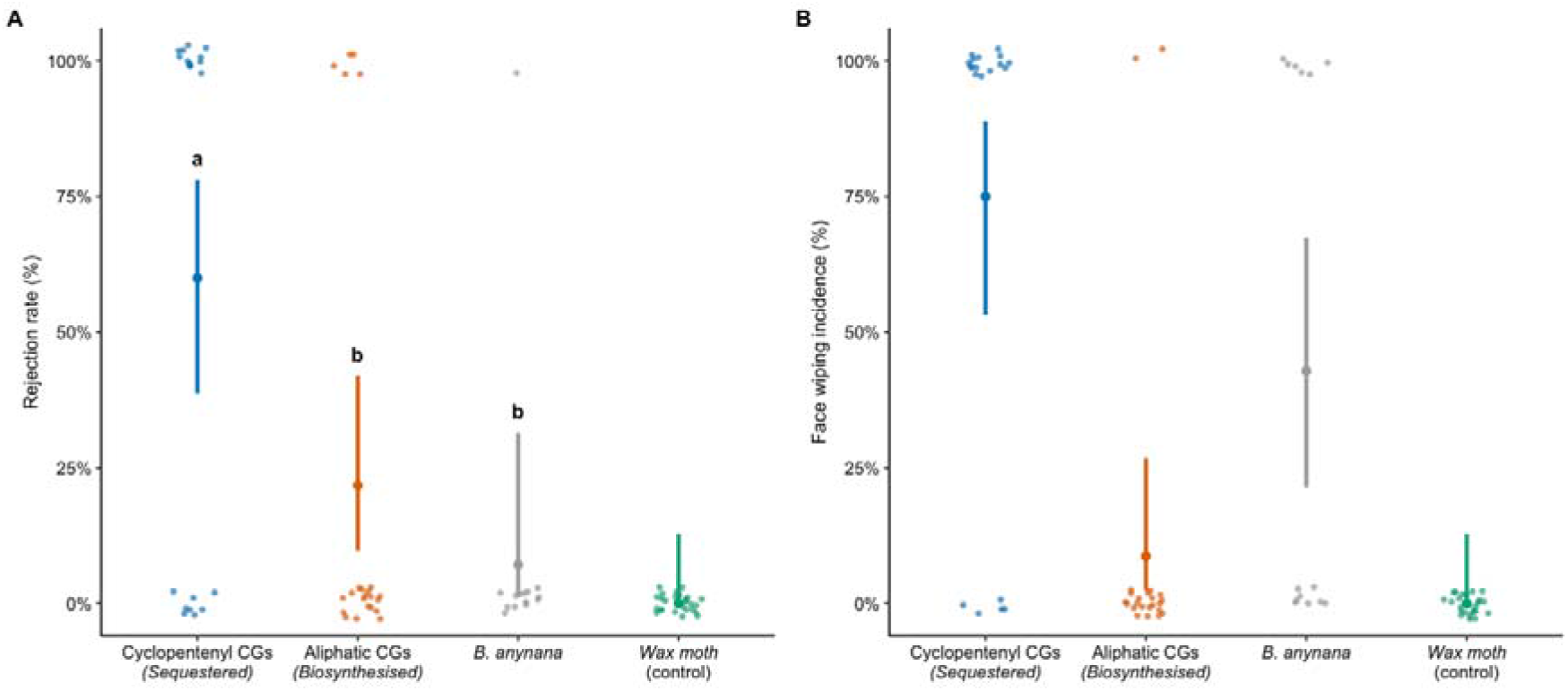
Rejection rate (a) and face wiping incidence (b) of Hierodula membranacea mantids presented with Heliconius sara larvae and palatable controls. (n = 20 cyclopentenyl CG, 23 aliphatic CG, 14 B. anynana, 26 control). Points show observed proportions; error bars show Wilson 95% CIs. Controls (0% rejection) were excluded from statistical models due to complete separation and are shown descriptively. Different letters denote significant pairwise differences (Tukey-adjusted, p < 0.05); no significant pairwise differences were found for face wiping (closest comparison p = 0.073).

Mantids that attacked *H. sara* larvae with cyclopentenyl CGs displayed face wiping at a substantially higher rate than the other groups, with 15 of 20 trials (75.0%) resulting in face wiping, compared with 2 of 23 *H. sara* with aliphatic CGs (8.7%), 6 of 14 *B. anynana* (42.9%) and 0 of 26 wax moth controls (0%) (Figure 1b). A binomial GLMM, including mantid identity as a random effect, indicated that larvae with cyclopentenyl CGs elicited face wiping more often than larvae with aliphatic CGs, although this difference did not reach significance after Tukey correction (estimate = 5.58, SE = 2.55, z = 2.191, p = 0.0726). Face wiping did not differ significantly between *H. sara* larvae with cyclopentenyl CGs and *B. anynana* (estimate = 2.74, SE = 1.52, z = 1.805, p = 0.168), or between *H. sara* larvae with aliphatic CGs and *B. anynana* (estimate = −2.84, SE = 1.56, z = −1.818, p = 0.164). Larval weight did not significantly predict face wiping (estimate = −2.09, SE = 1.388, z = −1.505, p = 0.132).

### Ants interact equally with butterfly larvae despite structural differences in their CG profile

We next explored a different invertebrate predator to determine the generality of our results with mantids. Ants spent significantly less total time interacting with larvae of both *H. sara* treatment groups than with wax moth controls (*H. sara* with cyclopentenyl CGs vs control: ratio = 0.321, t = −5.180, p < 0.0001; *H. sara* with aliphatic CGs vs control: ratio = 0.330, t = −4.735, p < 0.0001), with mean total interaction times of 36.8s (95% CI: 27.7–48.8s), 37.8s (95% CI: 27.3–52.4s), and 114.6s (95% CI: 82.2–159.8s) for *H. sara* with cyclopentenyl CG, larvae with aliphatic CGs, and control larvae respectively (Figure 2). Control larvae were significantly larger than both *H. sara* treatment groups (one-way ANOVA: F(2,86) = 179.7, p < 0.001), while the two *H. sara* groups did not differ significantly in size (p = 0.768); comparisons between experimental and control groups should be interpreted accordingly. Total interaction time did not differ significantly between *H. sara* treatment groups (ratio = 0.973, t = −0.126, p = 0.991).

**Figure 2:**
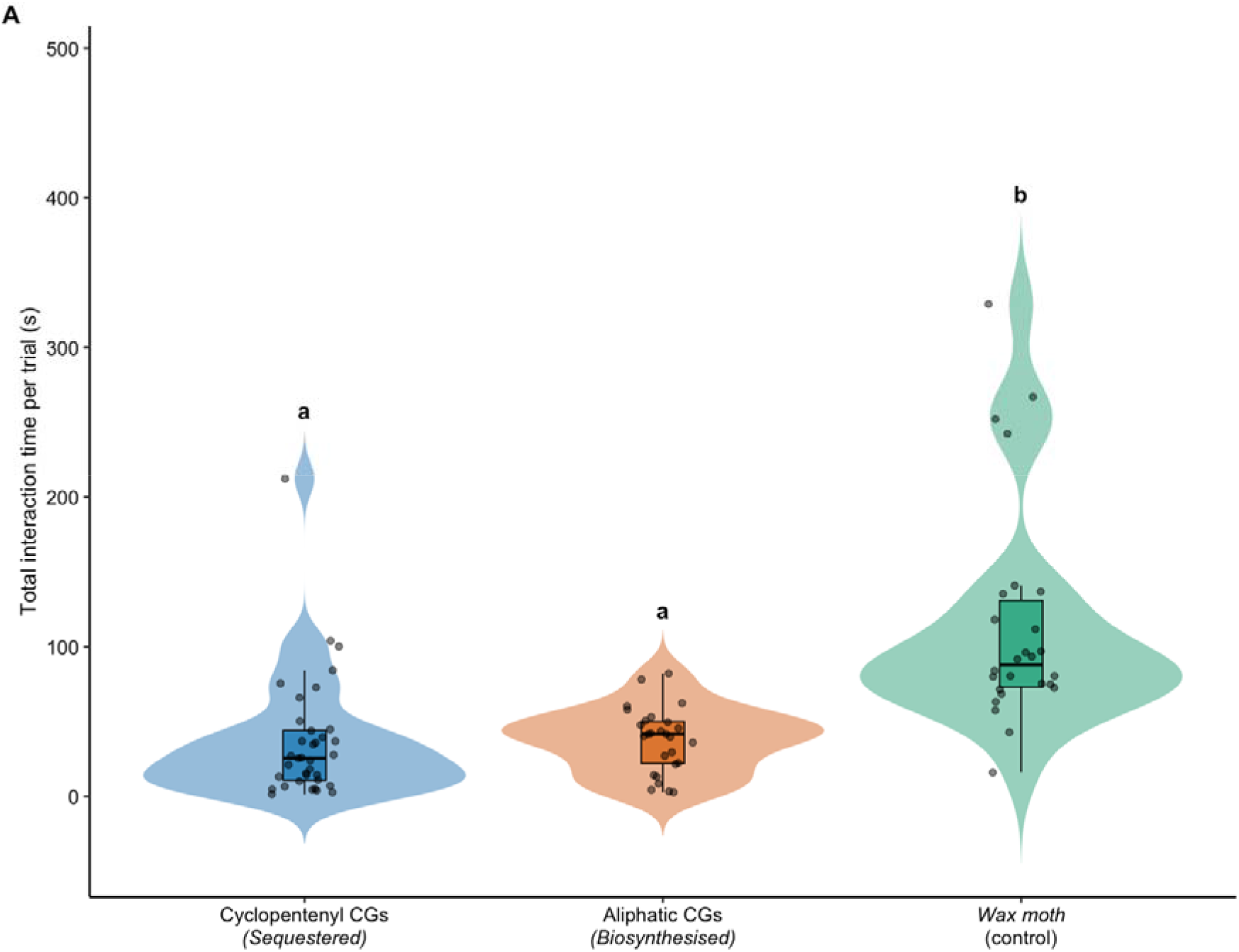
Total interaction time between Camponotus floridanus workers and intact Heliconius sara larvae raised on different host plants. (n = 36 cyclopentenyl CG, 27 aliphatic CG, 26 control). Violin plots show the full distribution of trial-level interaction times; boxplots show median and interquartile range. Different letters denote significant pairwise differences (Tukey-adjusted, p < 0.05). Control larvae were significantly larger than experimental larvae (F(2,86) = 179.7, p < 0.001); comparisons involving controls should be interpreted accordingly.

### Beyond toxicity: intact cyclopentenyl CGs are more unpalatable to ants than intact aliphatic CGs

In order to separate the effect of chemical structures from other differences between control and Heliconius larvae, such as defensive spines, we next applied chemical extracts directly to control larvae. When CG extracts were applied to wax moth larvae and presented to ants, CG structure significantly affected total interaction time. Ants interacted substantially less with wax moth larvae painted with extract containing cyclopentenyl CGs (sequestered) (mean: 107s, 95% CI: 85.6–133s) than with those painted with extract containing aliphatic CGs (biosynthesised) (mean: 215s, 95% CI: 154.9–298s) or water controls (mean: 242s, 95% CI: 182.3–320s) (extract with cyclopentenyl CGs vs extract with aliphatic CGs: ratio = 0.497, t = −2.766, p = 0.017; extract with cyclopentenyl CGs vs control: ratio = 0.441, t = −4.591, p < 0.0001) (Figure 3). Total interaction time did not differ significantly between larvae painted with extract with aliphatic CGs and controls (ratio = 0.888, t = −0.569, p = 0.837).

**Figure 3:**
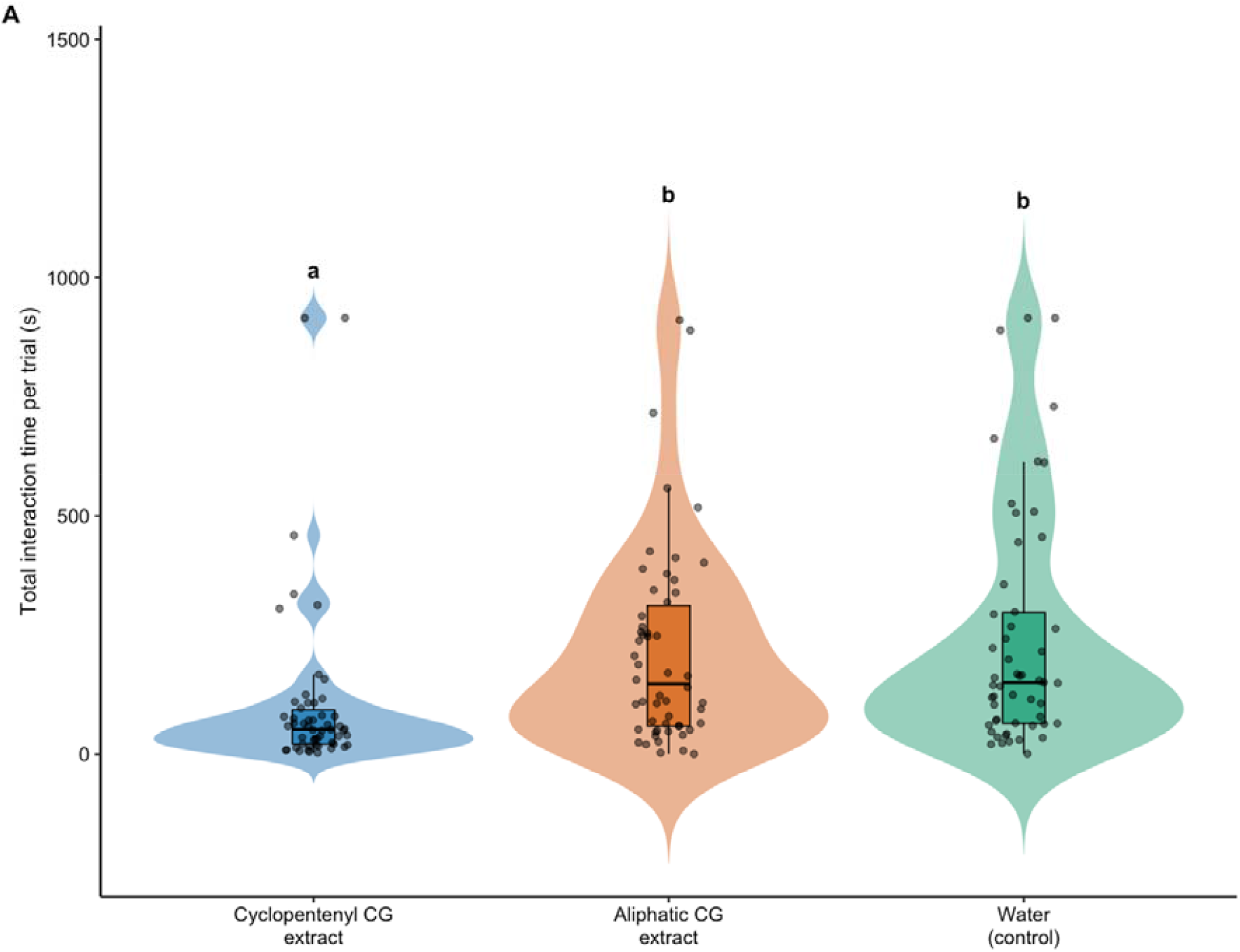
Total interaction time between Camponotus floridanus workers and wax moth larvae painted with cyclopentenyl or aliphatic cyanogenic glucoside extracts derived from H. sara, or water controls. (n = 54 per group). Violin plots show the full distribution of trial-level interaction times. Different letters denote significant pairwise differences (Tukey-adjusted, p < 0.05).

## Discussion

### Predation can act on unpalatability independently of toxicity

Our results demonstrate that sequestered cyclopentenyl CGs confer more protection to *Heliconius* butterflies against invertebrate predators than biosynthesised CGs. While predation has been shown to be impacted by toxicity [41], [42], [43], structural variation has received comparatively less attention [44] (but see [45]). Our findings emphasise that, even within the same class of toxins, distinct compounds can vary considerably in their palatability.

Mantids rejected *H. sara* larvae with cyclopentenyl CG (sequestered) more often than those with aliphatic CG (biosynthesised), despite comparable total CG content. Distress after prey consumption (face wiping) showed a similar, though marginally non-significant trend. Intact CGs are not toxic but are broken down into toxic cyanide and volatile aglycones upon tissue disruption [46]. As aliphatic and cyclopentenyl CGs release the same amount of cyanide, the repulsiveness of the latter to mantids must be due to the taste or smell of the compounds themselves or their volatile aglycones. This is consistent with previously demonstrated mantid responsiveness to olfactory cues from chemically defended prey [47]. Our results are also consistent with past research, where it has been shown that variation in the structure of breakdown products of aromatic CGs were more distasteful to chicks than those of aliphatic CGs despite comparable cyanide concentration [45]. As cyanide-release is not instantaneous, predators may be responding to cues associated with CG structure rather than to cyanide toxicity alone.

We disentangled unpalatability from toxicity by exposing ants to wax moth larvae painted with intact CG extracts from *H. sara*. Our extraction protocol disabled cyanogenesis, therefore eliminating the toxicity of CGs. Ants were more repulsed by larvae painted with cyclopentenyl CGs than aliphatic CGs, as shown through a reduced handling time. Indeed, in the absence of cyanogenesis, larvae painted with aliphatic CGs were no more repulsive than control larvae painted with water. To our knowledge, this is the first demonstration that intact toxin precursors are aversive to predators, extending previous findings that invertebrate predators can respond to non-toxic aspects of chemical defences independent of toxicity [45], [48]. Unpalatability might also act synergistically with toxicity to enhance predator learning of prey chemical defences. Indeed, an experiment using avian model predators found that unpalatability alone was insufficient to drive predator learning in sequential encounters which mimicked natural conditions [49].

### Evolutionary consequences of structural variation in chemical defences

Our results demonstrate that structural variation in chemical defences can have meaningful fitness implications. Other properties of a compound, in addition to toxicity, can shape its fitness consequences. Centipede venom has been shown to differentially impact conspecifics and heterospecifics, likely in an adaptive manner [50]. This shows that even subtle structural or target-specificity differences in a single compound can produce distinct outcomes for different recipients (also [51], [52], [53]), raising the possibility that distinct antagonists could select for different toxin structures within a species.

Our results also carry a related implication for quasi-Batesian mimicry, where co-mimics are typically assumed to differ mainly in toxin quantity [54]. We show that toxin structure alone can produce large differences in palatability even when toxin quantity is matched, meaning a co-mimic could be functionally under-defended even where toxin concentrations are similar.

Within *Heliconius*, biosynthesis maintains levels of toxicity independent of hostplant availability [12], [13]. However, our results suggest that sequestration produces a more deterrent chemical defence profile in some contexts. This could help explain why plasticity of toxin acquisition is maintained within many *Heliconius* species, giving an advantage to sequestration of plant compounds despite its potential costs in terms of increased dependency on plant defences [55]. Furthermore, in species where sequestration draws on multiple, chemically distinct host plants (e.g. [56], [57]), the resulting toxins may differ even more fundamentally than the CG classes compared here, potentially giving structure an even greater role in shaping unpalatability.

Taken together, our results suggest CG structure affects fitness benefits depending on the ecological context. Cyclopentenyl CGs are considerably less palatable to insect predators even when intact, as compared to the aliphatic compounds that *Heliconius* butterflies can synthesise themselves. More broadly, our results indicate that predation can impose selection pressures directly on the structure of the compounds, not only on their concentration. This demonstrates that unpalatability is important in the evolution of chemical defences and should be considered alongside overall toxicity.

## Supporting information

Supplementary statistical analysis

## Data accessibility

Data collected during trials can be found at [58], tracking code used to analyse filmed trials can be found at [32], and the code used to generate all statistics reported in this paper is at [59].

## Notes

https://zenodo.org/records/22641973

https://doi.org/10.5281/zenodo.22812978

https://doi.org/10.5281/zenodo.22811282

### Competing Interest Statement

The authors have declared no competing interest.

### Funding

- NERC Environmental Omics Facility: NE/W005131/1

## References

[1] N. M. Marples, M. P. Speed, and R. J. Thomas, ‘An individual-based profitability spectrum for understanding interactions between predators and their prey’, Biological Journal of the Linnean Society, vol. 125, no. 1. pp. 1–13, 2018.

[2] J. Mappes, N. Marples, and J. A. Endler, ‘The complex business of survival by aposematism’, Trends in Ecology & Evolution, vol. 20, no. 11, pp. 598–603, Nov. 2005, doi: 10.1016/j.tree.2005.07.011.

[3] M. Chouteau, M. Arias, and M. Joron, ‘Warning signals are under positive frequency-dependent selection in nature’, Proceedings of the National Academy of Sciences, vol. 113, no. 8, pp. 2164–2169, Feb. 2016, doi: 10.1073/pnas.1519216113.

[4] C. R. Darst, M. E. Cummings, and D. C. Cannatella, ‘A mechanism for diversity in warning signals: Conspicuousness versus toxicity in poison frogs’, Proceedings of the National Academy of Sciences, vol. 103, no. 15, pp. 5852–5857, Apr. 2006, doi: 10.1073/pnas.0600625103.

[5] E. S. Briolat et al., ‘Diversity in warning coloration: selective paradox or the norm?’, Biological Reviews, vol. 94, no. 2, pp. 388–414, Apr. 2019, doi: 10.1111/brv.12460.

[6] M. Chouteau, J. Dezeure, T. N. Sherratt, V. Llaurens, and M. Joron, ‘Similar predator aversion for natural prey with diverse toxicity levels’, Animal Behaviour, vol. 153, pp. 49–59, Jul. 2019, doi: 10.1016/j.anbehav.2019.04.017.

[7] M. P. Speed, G. D. Ruxton, J. Mappes, and T. N. Sherratt, ‘Why are defensive toxins so variable? An evolutionary perspective’, Biological Reviews, vol. 87, no. 4, pp. 874–884, Nov. 2012, doi: 10.1111/j.1469-185X.2012.00228.x.

[8] A. M. Winsor, M. Ihle, and L. A. Taylor, ‘Methods for independently manipulating palatability and color in small insect prey.’, PLoS One, vol. 15, no. 4, p. e0231205, 2020, doi: 10.1371/journal.pone.0231205.

[9] B. Rojas et al., ‘How to fight multiple enemies: target-specific chemical defences in an aposematic moth’, Proceedings of the Royal Society B: Biological Sciences, vol. 284, no. 1863, p. 20171424, Sep. 2017, doi: 10.1098/rspb.2017.1424.

[10] M. P. Speed, ‘Batesian, quasi-Batesian or Müllerian mimicry? Theory and data in mimicry Research’, Evolutionary Ecology, vol. 13, no. 7, pp. 755–776, Nov. 1999, doi: 10.1023/A:1010871106763.

[11] M. P. Speed, ‘Muellerian mimicry and the psychology of predation’, Animal Behaviour, vol. 45, no. 3, pp. 571–580, Mar. 1993, doi: 10.1006/anbe.1993.1067.

[12] D. Mebs, ‘Toxicity in animals. Trends in evolution?’, Toxicon, vol. 39, no. 1, pp. 87–96, Jan. 2001, doi: 10.1016/S0041-0101(00)00155-0.

[13] S. E. W. Opitz and C. Müller, ‘Plant chemistry and insect sequestration’, Chemoecology, vol. 19, no. 3, pp. 117–154, Sep. 2009, doi: 10.1007/s00049-009-0018-6.

[14] É. C. Pinheiro de Castro, M. Zagrobelny, J. P. Zurano, M. Zikan Cardoso, R. Feyereisen, and S. Bak, ‘Sequestration and biosynthesis of cyanogenic glucosides in passion vine butterflies and consequences for the diversification of their host plants’, Ecology and Evolution, vol. 9, no. 9, pp. 5079–5093, May 2019, doi: 10.1002/ece3.5062.

[15] F. Beran and G. Petschenka, ‘Sequestration of Plant Defense Compounds by Insects: From Mechanisms to Insect–Plant Coevolution’, Annual Review of Entomology, vol. 67, no. Volume 67, 2022. Annual Reviews, pp. 163–180, 2022. doi: 10.1146/annurev-ento-062821-062319.

[16] É. C. P. de Castro, J. Musgrove, S. Bak, W. O. McMillan, and C. D. Jiggins, ‘Phenotypic plasticity in chemical defence of butterflies allows usage of diverse host plants’, Biology Letters, vol. 17, no. 3, p. 20200863, Mar. 2021, doi: 10.1098/rsbl.2020.0863.

[17] A. L. K. Mattila, C. D. Jiggins, and M. Saastamoinen, ‘Condition dependence in biosynthesized chemical defenses of an aposematic and mimetic Heliconius butterfly’, Ecology and Evolution, vol. 12, no. 6, p. e9041, Jun. 2022, doi: 10.1002/ece3.9041.

[18] S. Pentzold, M. Zagrobelny, P. S. Roelsgaard, B. L. Møller, and S. Bak, ‘The Multiple Strategies of an Insect Herbivore to Overcome Plant Cyanogenic Glucoside Defence’, PLOS ONE, vol. 9, no. 3, p. e91337, Mar. 2014, doi: 10.1371/journal.pone.0091337.

[19] M. Zagrobelny, S. Bak, and B. L. Møller, ‘Cyanogenesis in plants and arthropods’, Phytochemistry, vol. 69, no. 7, pp. 1457–1468, May 2008, doi: 10.1016/j.phytochem.2008.02.019.

[20] M. E. Alonso-Amelot and A. Oliveros-Bastidas, ‘Kinetics of the Natural Evolution of Hydrogen Cyanide in Plants in Neotropical Pteridium arachnoideum and itsEcological Significance’, Journal of Chemical Ecology, vol. 31, no. 2, pp. 315–331, Feb. 2005, doi: 10.1007/s10886-005-1343-z.

[21] H. S. Engler, K. C. Spencer, and L. E. Gilbert, ‘Preventing cyanide release from leaves’, Nature, vol. 406, no. 6792, pp. 144–145, Jul. 2000, doi: 10.1038/35018159.

[22] P. Pokharel, M. Sippel, A. Vilcinskas, and G. Petschenka, ‘Defense of Milkweed Bugs (Heteroptera: Lygaeinae) against Predatory Lacewing Larvae Depends on Structural Differences of Sequestered Cardenolides’, Insects, vol. 11, no. 8, p. 485, 2020, doi: 10.3390/insects11080485.

[23] D. Mebs, C. Wunder, W. Pogoda, and S. W. Toennes, ‘Feeding on toxic prey. The praying mantis (Mantodea) as predator of poisonous butterfly and moth (Lepidoptera) caterpillars’, Toxicon, vol. 131, pp. 16–19, Jun. 2017, doi: 10.1016/j.toxicon.2017.03.010.

[24] P. Singh, N. Grone, L. J. Tewes, and C. Müller, ‘Chemical defense acquired via pharmacophagy can lead to protection from predation for conspecifics in a sawfly’, Proceedings of the Royal Society B: Biological Sciences, vol. 289, no. 1978, p. 20220176, Jul. 2022, doi: 10.1098/rspb.2022.0176.

[25] S. Mohammadi, L. Yang, M. Bulbert, and H. M. Rowland, ‘Defence mitigation by predators of chemically defended prey integrated over the predation sequence and across biological levels with a focus on cardiotonic steroids’, Royal Society Open Science, vol. 9, no. 9, p. 220363, Sep. 2022, doi: 10.1098/rsos.220363.

[26] L. P. M. Macêdo, E. O. Silva, and A. C. A. de Aguiar-Dias, ‘Morphoanatomy and ecology of the extrafloral nectaries in two species of Passiflora L. (Passifloraceae)’, South African Journal of Botany, vol. 143, pp. 248–255, Dec. 2021, doi: 10.1016/j.sajb.2021.07.040.

[27] J. Apple and D. Feener Jr, ‘Ant visitation of extrafloral nectaries of Passiflora: The effects of nectary attributes and ant behavior on patterns in facultative ant-plant mutualisms’, Oecologia, vol. 127, pp. 409–416, May 2001, doi: 10.1007/s004420000605.

[28] J. T. Smiley, ‘Heliconius Caterpillar Mortality during Establishment on Plants With and Without Attending Ants’, Ecology, vol. 66, no. 3, pp. 845–849, Jun. 1985, doi: 10.2307/1940546.

[29] J. M. G. Segovia and S. Pekár, ‘Aversive reactions of two invertebrate predators to European red–black insects’, Ethology, vol. 129, no. 1, pp. 24–32, Jan. 2023, doi: 10.1111/eth.13341.

[30] C. J. Paradise and N. E. Stamp, ‘Prey recognition time of praying mantids (Dictyoptera: Mantidae) and consequent survivorship of unpalatable prey (Hemiptera: Lygaeidae)’, Journal of Insect Behavior, vol. 4, no. 3, pp. 265–273, May 1991, doi: 10.1007/BF01048277.

[31] V. H. Sridhar, vivekhsridhar/tracktor: Tracktor. (Dec. 2017). Zenodo. doi: 10.5281/zenodo.1134016.

[32] E. Todd, ErikTodd12/Toxin-structure-shapes-palatability-in-a-chemically-defended-butterfly: Predator–prey interaction tracking pipeline for Heliconius sara palatability assays. (Sep. 2026). Zenodo. doi: 10.5281/zenodo.22641973.

[33] J. Cheng, Y. Zhou, K. Gu, Y. Zhang, and X. Feng, ‘Extraction and liquid chromatographic analysis of cyanogenic glycosides: a review’, Journal of Chromatography A, vol. 1760, p. 466292, Oct. 2025, doi: 10.1016/j.chroma.2025.466292.

[34] R Core Team, ‘R: A Language and Environment for Statistical Computing’. R Foundation for Statistical Computing, Vienna, Austria, 2026. [Online]. Available: https://www.R-project.org/

[35] F. Hartig, DHARMa: Residual Diagnostics for Hierarchical (Multi-Level / Mixed) Regression Models. 2024. [Online]. Available: https://CRAN.R-project.org/package=DHARMa

[36] R. V. Lenth and J. Piaskowski, emmeans: Estimated Marginal Means, aka Least-Squares Means. 2025. [Online]. Available: https://CRAN.R-project.org/package=emmeans

[37] M. E. Brooks et al., ‘glmmTMB Balances Speed and Flexibility Among Packages for Zero-inflated Generalized Linear Mixed Modeling’, The R Journal, vol. 9, no. 2, pp. 378–400, 2017, doi: 10.32614/RJ-2017-066.

[38] M. McGillycuddy, D. I. Warton, G. Popovic, and B. M. Bolker, ‘Parsimoniously Fitting Large Multivariate Random Effects in glmmTMB’, Journal of Statistical Software, vol. 112, no. 1, pp. 1–19, 2025, doi: 10.18637/jss.v112.i01.

[39] A. Zeileis, S. Köll, and N. Graham, ‘Various Versatile Variances: An Object-Oriented Implementation of Clustered Covariances in R’, J. Stat. Soft., vol. 95, no. 1, pp. 1–36, Oct. 2020, doi: 10.18637/jss.v095.i01.

[40] A. Zeileis and T. Hothorn, ‘Diagnostic Checking in Regression Relationships’, R News, vol. 2, no. 3, pp. 7–10, 2002.

[41] C. Barnett, J. Skelhorn, M. Bateson, and C. Rowe, ‘Educated predators make strategic decisions to eat defended prey according to their toxin content’, Behavioral Ecology, vol. 23, pp. 418–424, Dec. 2011, doi: 10.1093/beheco/arr206.

[42] R. A. Hayes, M. R. Crossland, M. Hagman, R. J. Capon, and R. Shine, ‘Ontogenetic Variation in the Chemical Defenses of Cane Toads (Bufo marinus): Toxin Profiles and Effects on Predators’, Journal of Chemical Ecology, vol. 35, no. 4, pp. 391–399, Apr. 2009, doi: 10.1007/s10886-009-9608-6.

[43] J. Skelhorn and C. Rowe, ‘Avian predators taste-reject aposematic prey on the basis of their chemical defence.’, Biol Lett, vol. 2, no. 3, pp. 348–350, Sep. 2006, doi: 10.1098/rsbl.2006.0483.

[44] M. P. Speed, G. D. Ruxton, J. Mappes, and T. N. Sherratt, ‘Why are defensive toxins so variable? An evolutionary perspective’, Biological Reviews, vol. 87, no. 4, pp. 874–884, 2012, doi: 10.1111/j.1469-185X.2012.00228.x.

[45] M. Z. Cardoso, ‘The effect of cyanogenic glucosides and their breakdown products on predation by domestic chicks’, Chemoecology, vol. 30, no. 3, pp. 131–138, Jun. 2020, doi: 10.1007/s00049-020-00304-6.

[46] M. Zagrobelny, É. C. De Castro, B. L. Møller, and S. Bak, ‘Cyanogenesis in Arthropods: From Chemical Warfare to Nuptial Gifts’, Insects, vol. 9, no. 2, p. 51, 2018, doi: 10.3390/insects9020051.

[47] L. E. Schweikert et al., ‘Experience with Aposematic Defense Triggers Attack Bias in a Mantid Predator (Stagmomantis carolina).’, Integr Org Biol, vol. 6, no. 1, p. obae039, 2024, doi: 10.1093/iob/obae039.

[48] S. C. Peterson, ‘Breakdown products of cyanogenesis’, Naturwissenschaften, vol. 73, no. 10, pp. 627–628, Oct. 1986, doi: 10.1007/BF00368782.

[49] J. Skelhorn and C. Rowe, ‘Distastefulness as an antipredator defence strategy’, Animal Behaviour, vol. 78, no. 3, pp. 761–766, Sep. 2009, doi: 10.1016/j.anbehav.2009.07.006.

[50] S. Yang et al., ‘Target switch of centipede toxins for antagonistic switch’, Science Advances, vol. 6, no. 32, p. eabb5734, doi: 10.1126/sciadv.abb5734.

[51] B. Panossian et al., ‘Phage toxin variants are linked to protection specificity in a defensive symbiont.’, Mol Biol Evol, vol. 43, no. 5, May 2026, doi: 10.1093/molbev/msag079.

[52] C. M. Modahl Mrinalini, S. Frietze, and S. P. Mackessy, ‘Adaptive evolution of distinct prey-specific toxin genes in rear-fanged snake venom.’, Proc Biol Sci, vol. 285, no. 1884, Aug. 2018, doi: 10.1098/rspb.2018.1003.

[53] O. Michálek, G. F. King, and S. Pekár, ‘Prey specificity of predatory venoms’, Biological Reviews, vol. 99, no. 6, pp. 2253–2273, Dec. 2024, doi: 10.1111/brv.13120.

[54] M. Arias et al., ‘Variation in cyanogenic compounds concentration within a Heliconius butterfly community: does mimicry explain everything?’, BMC Evol Biol, vol. 16, no. 1, p. 272, Dec. 2016, doi: 10.1186/s12862-016-0843-5.

[55] É. C. P. de Castro, J. Musgrove, S. Bak, W. O. McMillan, and C. D. Jiggins, ‘Phenotypic plasticity in chemical defence of butterflies allows usage of diverse host plants’, Biology Letters, vol. 17, no. 3, p. 20200863, Mar. 2021, doi: 10.1098/rsbl.2020.0863.

[56] L. Espinosa del Alba and G. Petschenka, ‘No physiological costs of dual sequestration of chemically different plant toxins in the milkweed bug Spilostethus saxatilis (Heteroptera: Lygaeidae)’, Journal of Insect Physiology, vol. 147, p. 104508, Jun. 2023, doi: 10.1016/j.jinsphys.2023.104508.

[57] C. C. M. Arce et al., ‘The polyvalent sequestration ability of an economically important beetle’, Current Biology, vol. 34, no. 23, pp. 5417–5428.e4, Dec. 2024, doi: 10.1016/j.cub.2024.10.005.

[58] E. Todd and A. Denton, ‘Toxin structure shapes palatability in a chemically defended butterfly - Data’. Zenodo, Sep. 17, 2026. doi: 10.5281/ZENODO.22811282.

[59] E. Todd, ErikTodd12/Toxin-structure-shapes-palatability-in-a-chemically-defended-butterfly—statistics: Statistics for the paper: Toxin structure shapes palatability in a chemically defended butterfly. (Sep. 2026). Zenodo. doi: 10.5281/zenodo.22812978.

