## Supplementary statistical analysis for "Toxin structure shapes palatability in a chemically defended butterfly"

This document reports the full statistical output underlying the results presented in the main text, including model summaries, pairwise contrasts, and model validity checks for each of the three experiments. All analyses were conducted in R (version 4.3.3), using the packages and model structures described in the Methods.

The code used to produce all of the statistics below can be found at <https://doi.org/10.5281/zenodo.22812978>. All data has been deposited at [10.5281/zenodo.22811282](https://doi.org/10.5281/zenodo.22811282).

### Experiment 1: Ant assay, live larvae

Total interaction time between *C. floridanus* workers and live *H. sara* larvae (cyclopentenyl CG, aliphatic CG) or wax moth controls was modelled using a gamma GLM with a log link. No batch term was included, as larval availability meant treatment groups were not tested concurrently, and batch and treatment were confounded.

#### Gamma GLM: total interaction time

| **Term** | **Estimate** | **SE** | **t value** | **p value** |
| --- | --- | --- | --- | --- |
| (Intercept) | 3.605 | 0.142 | 25.365 | < 2e-16 |
| Biflora | 0.027 | 0.217 | 0.126 | 0.900 |
| Control | 1.137 | 0.219 | 5.180 | 1.44e-06 |

Dispersion parameter (Gamma family): 0.727. Null deviance: 89.253 on 88 df. Residual deviance: 63.225 on 86 df. AIC: 879.54.

#### Pairwise contrasts (Tukey-adjusted, back-transformed)

| **Contrast** | **Ratio** | **SE** | **df** | **t ratio** | **p value** |
| --- | --- | --- | --- | --- | --- |
| Auriculata / Biflora | 0.973 | 0.211 | 86 | -0.126 | 0.991 |
| Auriculata / Control | 0.321 | 0.070 | 86 | -5.180 | < 0.0001 |
| Biflora / Control | 0.330 | 0.077 | 86 | -4.735 | < 0.0001 |

Estimated marginal means (back-transformed, seconds): Auriculata 36.8 (95% CI: 27.7-48.8), Biflora 37.8 (95% CI: 27.3-52.4), Control 114.6 (95% CI: 82.2-159.8).

#### Larval size (weight)

Control larvae were confirmed to be significantly heavier than both *H. sara* treatment groups. A one-way ANOVA was fitted to larval weight by group, followed by Tukey pairwise comparisons.

| **Term** | **Df** | **Sum Sq** | **Mean Sq** | **F value** | **p value** |
| --- | --- | --- | --- | --- | --- |
| group | 2 | 0.920 | 0.460 | 179.7 | < 2e-16 |
| Residuals | 86 | 0.220 | 0.003 |  |  |

| **Contrast** | **Diff** | **Lower CI** | **Upper CI** | **p adj** |
| --- | --- | --- | --- | --- |
| Biflora - Auriculata | -0.009 | -0.040 | 0.022 | 0.768 |
| Control - Auriculata | 0.220 | 0.189 | 0.251 | < 0.0001 |
| Control - Biflora | 0.229 | 0.195 | 0.262 | < 0.0001 |

#### Model validity checks

| **Test** | **Statistic** | **Result** |
| --- | --- | --- |
| KS test (uniformity) | p = 0.685 | n.s. |
| Dispersion test | p = 0.688 | n.s. |
| Outlier test | p = 0.509 | n.s. |
| Within-group deviation from uniformity | - | n.s. |
| Levene test (homogeneity of variance) | - | n.s. |

*All checks for this model were clean*.

### Experiment 2: Ant assay, extract painting

Total interaction time between *C. floridanus* workers and wax moth larvae painted with cyanogenic glucoside extracts (cyclopentenyl, aliphatic) or water was modelled using a gamma GLM with a log link. A batch term (colony by day) was tested before being excluded from the final model.

#### Batch effect test

| **Model** | **Resid. df** | **Resid. dev** | **Df** | **Deviance** | **F** | **p value** |
| --- | --- | --- | --- | --- | --- | --- |
| group only | 159 | 212.35 |  |  |  |  |
| group + batch | 155 | 208.80 | 4 | 3.549 | 0.532 | 0.712 |

The batch term was non-significant and excluded from the final model. Robust standard errors clustered by batch were retained regardless as a conservative measure, and all reported estimates and pairwise comparisons for this experiment use these batch-clustered SEs.

#### Gamma GLM: total interaction time (model-based SEs)

| **Term** | **Estimate** | **SE** | **t value** | **p value** |
| --- | --- | --- | --- | --- |
| (Intercept) | 4.670 | 0.175 | 26.73 | < 2e-16 |
| Biflora | 0.699 | 0.247 | 2.83 | 0.00526 |
| Control | 0.818 | 0.247 | 3.31 | 0.00115 |

Dispersion parameter (Gamma family): 1.648. Null deviance: 231.53 on 161 df. Residual deviance: 212.35 on 159 df. AIC: 2010.

#### Coefficients with batch-clustered robust SEs

| **Term** | **Estimate** | **SE** | **z value** | **p value** |
| --- | --- | --- | --- | --- |
| (Intercept) | 4.670 | 0.111 | 41.931 | < 2.2e-16 |
| Biflora | 0.699 | 0.253 | 2.766 | 0.00567 |
| Control | 0.818 | 0.178 | 4.591 | 4.42e-06 |

#### Pairwise contrasts (Tukey-adjusted, robust SEs, back-transformed)

| **Contrast** | **Ratio** | **SE** | **df** | **t ratio** | **p value** |
| --- | --- | --- | --- | --- | --- |
| Auriculata / Biflora | 0.497 | 0.126 | 159 | -2.766 | 0.0173 |
| Auriculata / Control | 0.441 | 0.079 | 159 | -4.591 | < 0.0001 |
| Biflora / Control | 0.888 | 0.185 | 159 | -0.569 | 0.8369 |

Estimated marginal means (back-transformed, seconds): Auriculata 107 (95% CI: 85.6-133), Biflora 215 (95% CI: 154.9-298), Control 242 (95% CI: 182.3-320).

#### Model validity checks

| **Test** | **Statistic** | **Result** |
| --- | --- | --- |
| KS test (uniformity) | p = 0.142 | n.s. |
| Dispersion test | p = 0.832 | n.s. |
| Outlier test | p = 0.370 | n.s. |
| Within-group deviation from uniformity | - | n.s. |
| Levene test (homogeneity of variance) | - | n.s. |

All checks for this model were clean.

### Mantid experiment

Individual larvae (cyclopentenyl CG, aliphatic CG, *B. anynana*, wax moth control) were presented to single *H. membranacea* mantids. Wax moth controls were excluded from both models due to complete separation, as they produced zero events in both response variables.

#### Rejection

Rejection was modelled with a binomial GLM. Mantid identity was first tested as a random effect in a GLMM as a sanity check; its variance was negligible (approximately 4e-09), so it was dropped, and the plain GLM below is the final model used for reporting.

| **Contrast** | **Estimate** | **SE** | **z value** | **p value** |
| --- | --- | --- | --- | --- |
| Cyclopentenyl vs Aliphatic | 1.87 | 0.712 | 2.624 | 0.0236 |
| Cyclopentenyl vs B. anynana | 3.24 | 1.150 | 2.809 | 0.0138 |
| Aliphatic vs B. anynana | 1.37 | 1.160 | 1.184 | 0.463 |

Larval weight, included as a scaled covariate, was not a significant predictor of rejection.

##### Model validity checks

| **Test** | **Statistic** | **Result** |
| --- | --- | --- |
| KS test (uniformity) | p = 0.974 | n.s. |
| Dispersion test | dispersion = 1.031, p = 0.868 | n.s. |
| Outlier test | 0 outliers of 57, p = 1 | n.s. |
| Combined quantile test (qgam) | p = 0.753 | n.s. |
| Levene test, residuals by Type | - | n.s. |

#### Face wiping

Face wiping was modelled with a binomial GLMM including mantid identity as a random effect, since its variance (approximately 2.0) was clearly non-negligible. A naive GLM without this random effect was also fitted for comparison only; it is not used for reporting, as treating repeated trials from the same mantid as independent inflates significance through pseudoreplication. Under the naive model, the cyclopentenyl versus aliphatic contrast reached p = 0.0002; under the GLMM below, the same contrast is p = 0.0726.

| **Term** | **Estimate** | **SE** | **z value** | **p value** |
| --- | --- | --- | --- | --- |
| (Intercept) | 0.937 | 1.215 | 0.771 | 0.441 |
| Aliphatic | -5.580 | 2.546 | -2.191 | 0.0284 |
| *B. anynana* | -2.736 | 1.516 | -1.805 | 0.0711 |
| Larval weight (scaled) | -2.090 | 1.388 | -1.505 | 0.132 |

Random effect variance (mantid identity): 2.047 (SD = 1.431).

##### Pairwise contrasts (Tukey-adjusted, log-odds scale)

| **Contrast** | **Estimate** | **SE** | **df** | **z ratio** | **p value** |
| --- | --- | --- | --- | --- | --- |
| Cyclopentenyl - Aliphatic | 5.58 | 2.55 | Inf | 2.191 | 0.0726 |
| Cyclopentenyl - *B. anynana* | 2.74 | 1.52 | Inf | 1.805 | 0.1679 |
| Aliphatic - *B. anynana* | -2.84 | 1.56 | Inf | -1.818 | 0.1636 |

##### Model validity checks

| **Test** | **Statistic** | **Result** |
| --- | --- | --- |
| KS test (uniformity) | p = 0.608 | n.s. |
| Dispersion test | p = 0.646 | n.s. |
| Outlier test | p = 1 | n.s. |
| Zero-inflation test | p = 0.972 | n.s. |
| DHARMa residual vs predicted | - | no significant problems detected |

All checks for this model were clean.

### Overall summary

All four models pass their core DHARMa diagnostics (uniformity, dispersion, outlier, and homogeneity-of-variance tests), with no caveats to note.
